# Bioluminscent *Mycobacterium ulcerans*, a tool to study host-pathogen interactions in a murine tail model of Buruli ulcer

**DOI:** 10.1101/434506

**Authors:** Till F. Omansen, Renee A. Marcsisin, Brendon Y. Chua, Weiguang Zeng, David C. Jackson, Jessica L. Porter, Ymkje Stienstra, Tjip S. van der Werf, Timothy P. Stinear

**Author notes:** Corresponding author: Prof. Timothy P. Stinear, Department of Microbiology and Immunology, The Peter Doherty Institute for Infection and Immunity, University of Melbourne, Parkville, VIC 3010, Australia.

## Abstract

Buruli ulcer is a neglected tropical disease caused by infection with *Mycobacterium ulcerans*. In this study we used a previously reported strain of *M. ulcerans*, genetically engineered to constitutively produce bioluminescence, to follow the progression of Buruli ulcer in mice using an in-vivo imaging (IVIS^®^) system. We aimed to characterize a mouse tail infection model for pathogenesis, as well as for pre-clinical vaccine and drug development research for Buruli ulcer. Immune parameters, such as antibody titers and cytokine levels, were determined throughout the course of the infection and histology specimens were examined for comparison with human pathology. Nine out of ten (90%) BALB/c mice infected subcutaneously with 10^5^ *M. ulcerans* JKD8049 (containing pMV306 hsp16+luxG13) exhibited light emission from the site of infection over the course of the experiment indicating *M. ulcerans* growth *in-vivo*. Five out of ten (50%) animals developed clinical signs of disease. Antibody titers were overall low and their onset was late, as measured by responses to both heterogenous (bacterial whole cell lysate) and single antigen (Hsp18) targets. IFN-γ, and IL-10 are reported to play a vital role in host control of Buruli ulcer and these cytokines were elevated in animals with pathology. For mice with advanced pathology, histology revealed clusters of acid-fast bacilli within subcutaneous tissue 300-400 μm beneath the epidermis of the tail, with macrophage infiltration and granuloma-formation resembling human Buruli ulcer. This study has shown the utility of using bioluminescent *M. ulcerans* and IVIS^®^ in a mouse tail infection model to study Buruli ulcer infection.

**Author summary:** Buruli ulcer is one of the so called neglected tropical diseases. It is an infectious disease, mainly occurring in West Africa but also in Australia. It manifests as skin lesion and ulcer. Up to date, the way of transmission is inadequately understood. Also, there is no vaccine to protect against the disease. Buruli ulcer is treatable with a course of antibiotics that need to be given for the duration of two months. More laboratory research is needed to elucidate the mechanism of transmission, develop a vaccine and improve and shorten antibiotic therapy. For this, animal (mouse) models of disease are used. The aim of this study was to refine and improve the mouse tail infection model of Buruli ulcer. For this, we used a genetically modified *Mycobacterium ulcerans* strain that emits light. After infection of animals, light emitted from the bacteria was read out with an in-vivo imaging (IVIS) camera. This allowed us to monitor the location of bacteria in the living animal over time without the need to kill the animal. We also measured parameters of the immune system such as antibodies and cytokines as a baseline for future studies into immunology, vaccine development and pathology of Buruli ulcer. We successfully improved and characterized the mouse tail infection model in Buruli ulcer with the use of modern technology using light emitting bacteria and the IVIS camera.

## Introduction

*Mycobacterium ulcerans* causes the neglected tropical disease (NTD) Buruli ulcer (BU) that can manifest as a skin nodule, plaque, edematous lesion or open skin ulcer characterized by yellowish-white necrosis and undermined edges [1]. The disease generally occurs in clustered foci in rural Central and Western Africa but has also gained prominence in specific regions of south east Australia. Currently, 12 countries actively report Buruli ulcer cases and 33 have ever reported cases [2]. Patients with Buruli ulcer suffer from stigmatization, social participation restrictions and physical disability long after even treatment is complete [3]. The main pathogenic factor in BU is a diffusible cytotoxin called mycolactone. Mycolactone is a polyketide-derived macrolide that is responsible for the pathological triad of necrosis, suppressed local inflammatory response and hypoalgesia of the lesion [4,5]. By means of preventing protein translocation into the endoplasmic reticulum, mycolactone suppresses an efficient host innate and adaptive immune response [6,7]. The 174kb large plasmid pMUM001 is responsible for ML production by *M. ulcerans* [8].

There are several major challenges to control Buruli ulcer. The mode of transmission is not yet completely understood and seems to vary by geographic location, although puncturing injuries after contamination from an environmental source seem to be a major cause and in south east Australia at least, mosquitoes have been linked to transmission [9]. Buruli ulcer is currently treated with an eight-week regimen of rifampin and streptomycin or a regimen where the injectable streptomycin is replaced with clarithromycin after four weeks; a fully oral, eight-week rifampin and clarithromycin regimen has been trialled in humans and current trial analysis is ongoing (ClinicalTrials.gov Identifier: NCT01659437) [10,11]. Progressed, larger lesions are often managed with surgical excision of the infected tissue followed by functional repair and skin grafting [12]; a recent study showed that the time-point for decision making on whether to intervene surgically or not does not matter for overall healing outcomes [13]. No vaccine is available despite several efforts to employ the BCG-vaccine or to develop novel vaccines [14–23].

More preclinical research in *M. ulcerans* transmission, chemotherapy and vaccination as well as pathogenesis, is necessary to solve the biomedical challenges complicating Buruli ulcer infection control in the field. In this respect, *M. ulcerans* mouse-infection models have been pivotal in guiding research and clinical studies regarding these questions in the past [24–26]. Murine footpad and tail infection are established methods to study *M. ulcerans* [24,27]. The footpad-model has been derived from experience with experimental infection of *M. leprae* in mice [24,28]. It has been used in numerous pre-clinical studies, to mainly evaluate drug efficacy, but also vaccines for *M. ulcerans* [25,26,29–41]. Tail infection has been used to study pathology and vector research [27] and vaccinology [14]. Given that Buruli ulcer in humans is a subcutaneous infection mostly occurring on the lower and upper limbs [1,42], the mouse foot and tail are obvious sites to model the disease and the absence of fur in mice allows for easy clinical observation at these sites. Tail infection offers a cutaneous infection site that is not in contact with the environment as much as the footpad so that contamination, re-distribution or loss of inoculum and animal impairment in more advanced stages of the disease are less likely to occur. Also, it is a more practical region for imaging than the footpad. Bioluminescent strains of *M. ulcerans* have successfully been employed to evaluate drug efficacy in *in-vitro* and *in-vivo* drug efficacy studies [36,37,41,43] and in vector ecology studies of *M. ulcerans* [44]. Drug efficacy studies used a luminometer to assess light emission from infected mouse footpads [36,41,45].

In this study, we aimed to employed the use of a bioluminescent reporter strain of *M. ulcerans* to infect BALB/c mice and read out light emission using an in-vivo imaging system (IVIS^®^) that is both very sensitive at detecting light and enables us to visualize signals from the entire mouse body and thus localize the bacteria. This allows to confirm the site of infection and study possible spread of bacteria. We also characterized the immune-responses to this *M. ulcerans* strain in infected BALB/c mice over a long-term infection period of 17 weeks to establish a baseline for future transmission, vaccine and chemotherapy studies.

The bioluminescent *M. ulcerans* strain used in this study has been previously described and contains the pMV306 hsp16+luxG13 reporter plasmid [43,46,47] that integrates into the mycobacterial chromosome and contains the lux operon (luxABCDE). Thus, it does not require the addition of an exogenous substrate to detect bioluminescence [46]. Bioluminescent infection-models offer the possibility of *in vivo* imaging, with the luminescence read-out serving as proxy for bacterial burden. This approach greatly reduces mouse experiment sample sizes, preventing multiple animals to be killed for a common microbiological assessment such as counting bacterial colony-forming units (CFU) at different time points. It also offers a way to visualize disease progression over time, as well as bacterial spread in the living host.

## Materials and Methods

### Culture conditions

*M. ulcerans* JKD8049 harbouring pMV306 hsp16+luxG13 was grown on Middlebrook 7H10 agar or in 7H9 broth containing 10% Oleic Albumin Dextrose Catalase Growth Supplement (Middlebrook, Becton Dickinson, Sparks, MD, USA), 0.5% glycerol and 25μg/ml kanamycin sulfate (Amresco, Solon, OH, USA). Plates and flasks were incubated for 8-10 weeks at 30°C, 5% CO_2_. LC-MS was used to confirm that bioluminescent bacteria were still producing mycolactones [48].

### Establishing a standard curve for bioluminescent *M. ulcerans* JKD8049

Light emission in photos/sec was compared with colony-forming units (CFU) for *M. ulcerans* JKD8049 cultured in Middlebrook 7H9 medium for 4 weeks and then diluted in serial 10-fold steps in 96-well trays. Photon emissions were captured using a Lumina XRMS Series III In Vitro Imaging System (IVIS^®^) (Perkin Elmer, Waltham, MA, USA). Bacterial CFUs were confirmed by the spot plate method [9].

### Mouse-tail infections

Animal experimentation adhered to the Australian National Health and Medical Research Council Code for the Care and Use of Animals for Scientific Purposes and was approved by and performed in accordance with the University of Melbourne animal ethics committee (Application: 1312756.1). The animals were purchased from ARC (Canning Vale, Australia). Upon arrival, animals acclimatized for 5 days. Food and water were given *ad libitum*. Ten six-week old, female BALB/c mice were inoculated with approximately 10^5^ *M. ulcerans* CFU by subcutaneous (SC) injection into the dorsal aspect of the upper third of the tail. The concentration of the bacterial inocula was confirmed by spot plating. After 17 weeks post-inoculation or whenever the humane endpoint was reached, mice were humanely killed.

### *In-vivo* imaging

Mice were imaged once a week during morning time using a Lumina XRMS Series III IVIS^®^ as described above. During imaging, mice were anaesthetized with 2.5% isofluorane gas (Ceva Animal Health, Glenorie, NSW, Australia) with the stage on which the mice were placed during imaging was warmed to 37°C. Photon emissions were acquired with the following settings: exposure time 5 minutes, emission filter: open, excitation filter: blocked, binning: medium, F/stop 1. These images were superposed onto conventional black/white photographs (exposure time: auto, binning: medium, F/stop: 16). Images from the ventral and dorsal aspect of the tail were taken. Images were analyzed using Living Image^®^ software. Areas emitting light were defined as regions of interest (ROI). A copy of every ROI was placed next to those areas for background measurement. Photons per second from the ROIs were computed and values from control regions subtracted from the actual ROI. Results from ventral and dorsal images (Fig. 1B) were added and the cumulative luminescence of the two imaging angles reported.

**Figure 1.**
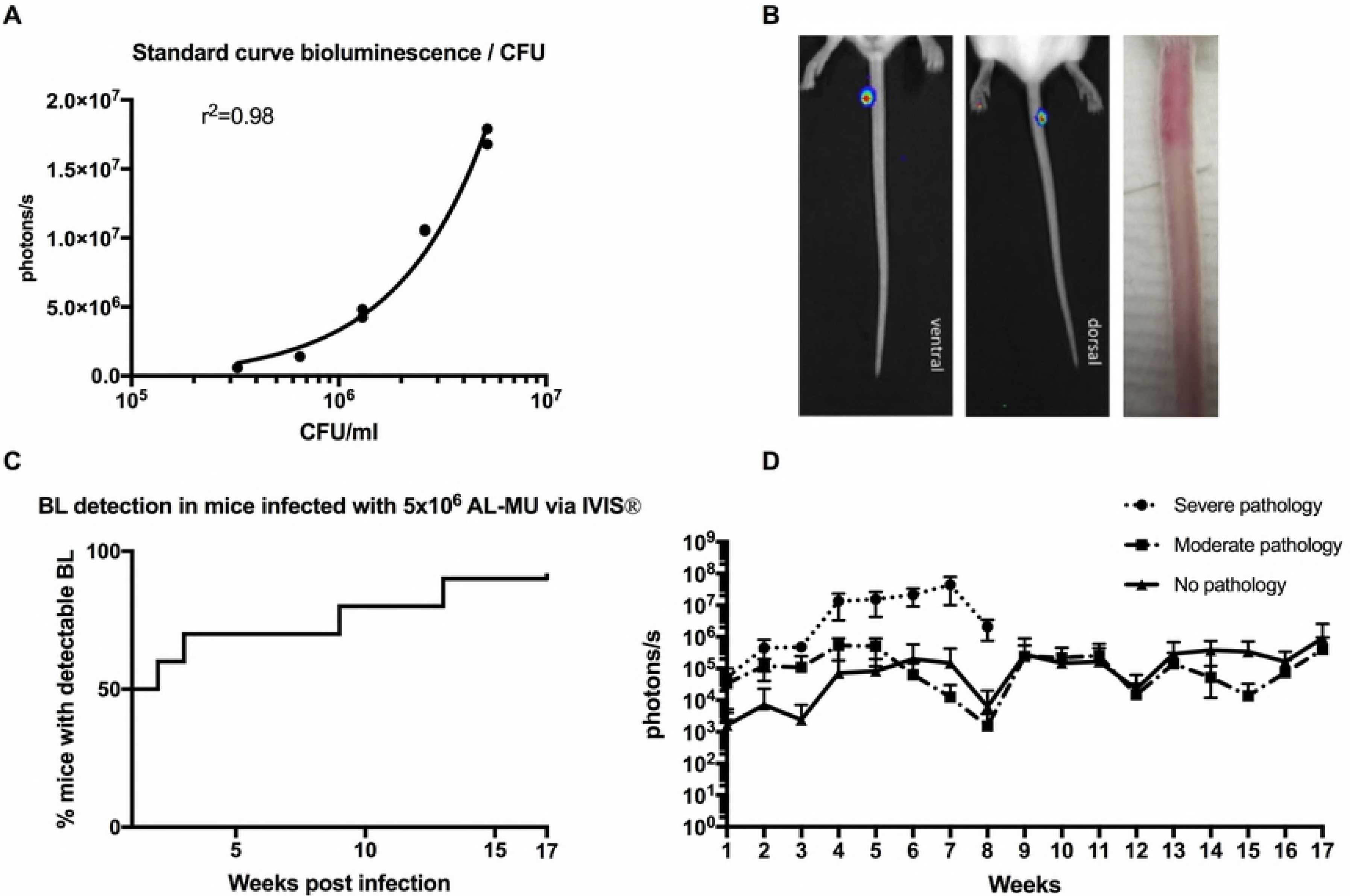
Use of bioluminescent *M. ulcerans* to follow the evolution of disease in the mouse tail model of Buruli ulcer. (A) Standard curve comparing bioluminescence and colony-forming units (CFU) of the *M. ulcerans* JKD8049 +pMV306 hsp16 luxG13 reporter strain. (B) IVIS^®^- (left) and photographic images (right) from BALB/c mice infected via subcutaneous tail inoculation with approx. 3.3×10^5^ CFU/ml *M. ulcerans* harboring the bioluminescent reporter plasmid. Photons/s emitted from bioluminescent bacteria were detected by IVIS^®^ in anesthetized mice. Results are represented in a pseudo-colored scheme (red indicated high, yellow medium and green low intensity of light emitted). Light was detected from both the dorsal (site of injection) and ventral aspects of the mouse tail. The photos were taken at 4 weeks post infection; the bioluminescence read-out was 1.8×10^6^ photons, corresponding to approx. 6.7×10^5^ CFU/ml according to our standard curve. (C) Survival-graph representing time to detectable bioluminescence emission from mice infected with bioluminescent *M. ulcerans* into the tail (D) Development of mean photons/s emitted from mice infected with 3.3×10^5^ bioluminescent *M. ulcerans* into the upper third of the dorsal tail. Values represent the mean of dorsal and ventral photons/s measurement. Animals were sub-grouped for analysis by clinical staging based on severity of the gross pathology (*severe*: redness, swelling, edema, impending ulceration; *moderate*: redness, edema; and no pathology.

### ELISA

Blood samples were obtained by sub-mandibular puncture and, at experimental endpoint, by cardiac puncture. Serum was collected by centrifugation and stored at −20°C. All incubation of ELISA-plates was done in a moisturized container at room temperature. Flat-bottom polyvinyl chloride microtiter-plates (Thermo Fischer, Milford, MA, USA) were coated with 5μg of the antigen overnight. Antigens used were the mycobacterial small heat-shock protein 18 (Hsp18) and *M. ulcerans* whole cell lysate (WCL). The antigen was discarded and plates blocked for 1h with PBS containing 10mg/ml bovine serum albumin (BSA). Plates were washed four times with PBS containing 0.05% Tween-20 (PBST). Sera were added in eight serial dilutions in PBS to the plate and incubated for 4 h. Plates were washed with PBST again and 50μl/well HRP-conjugated polyclonal rabbit anti-mouse Ig-antibody (Dako, Glostrup, Denmark) was added for 1h. Subsequently, ELISA substrate (0.2 mM 2,29-azino-bis 3-ethylbenzthiazoline-sulfonic acid in 50 mM citric acid containing 0.004% hydrogen peroxide) was added to detect bound antibodies. Absorbance was read in a plate reader at 405nm and 450nm and the average of the two wavelengths recorded.

### Intracellular cytokine staining and FACS

Dissected spleens were homogenized with a mesh and ATC treated. Splenocytes (1×10^6^) were re-stimulated with 2μg *M. ulcerans* JKD8049 whole cell lysate (WCL) in RPMI 1640 supplemented with 64mM L-glutamine, 32 mM sodium pyruvate, 1.75mM 2-mercaptoethanol, 3165μg/ml penicillin (all Gibco^®^ Life Technologies, NY, USA), 760 μg/ml gentamicin (G-Bioscience, St. Louis, MO, USA) and 10% heat-inactivated fetal calf serum (CSL, Parkville, Australia) for 72h at 37°C, 5% CO_2_. Plates were spun down, supernatant collected and stored at −20°C. Cytokines were stained using the bead-based Cytometric Bead Array (CBA) Mouse Th1/Th2/Th17 Cytokine Kit (BD, North Ryde, NSW, Australia) according to the manufacturer’s instructions. Samples were run on a BD FACSCanto™ II Flow Cytometry System and data analyzed using FCAP Array™ Analysis Software version 3.0.

### Histology

A section ranging approximately 5mm from the midline of the ulcer proximally was dissected and stored in 10% buffered formalin for histological assessment. Prepared paraffin blocks were surface-decalcified with 10% nitric acid for 5 minutes before cutting 4μm sections. Hematoxylin and eosin (HE) and Ziehl-Neelsen (ZN) straining were used following standard protocols. The specimens were subjected for analysis by an independent pathologist. Presence of AFB, inflammatory cells (macrophages, plasma cells/lymphocytes, neutrophils and eosinophils) as well as the degree of inflammation (granulomas, panniculitis, calcification, vasculitis, neuritis) and the tissue damage (dermal and fat tissue necrosis, muscle layer involvement and bone change) and the vascular involvement were scored. Specimens from two non-infected, naïve mice were used as controls.

### Statistical analysis

Statistical analysis was performed using GraphPad Prism version 7.0a (GraphPad Software, Inc., San Diego, CA). RLU data were graphed as mean of the ventral and dorsal reading, as described above. Time to bioluminescence is displayed as survival curve. Antibody titers are represented as the reciprocal of the highest dilutions of serum needed to measure an absorbance value of 0.2. This was achieved by transformation of the data by plotting absorbance values vs log0.5—fold dilutions data of each group and using a nonlinear regression analysis of to obtain a line of best fit (with 95% CI) to which the intersect value of 0.2 was determined (Prism 5 or whatever version). One-way ANOVA followed by a Tukey’s multiple comparisons test assuming an alpha of 0.05 was used to test for statistical significant difference between antibody titer measurements. Cytokine readings are shown compared using descriptive statistics.

## Results

### Standard curve comparing photon/s with CFU readout

To compare bioluminescence read-out with actual bacterial burden, we first established a standard curve *in-vitro*. We applied IVIS^®^ imaging to samples of JKD8049 pMV306 hsp16+luxG13 in different dilutions and plated these on agar for CFU determination. We were able to interpolate a standard curve by nonlinear regression showing a very high positive correlation (r^2^= 0.98) between photons/s and CFU/ml (Fig 1A).

### Establishment of mouse tail infection

In order to evaluate virulence and to study murine infection, bioluminescent *M. ulcerans* was injected subcutaneously into mouse tails. Ten BALB/c mice were inoculated with 3.3×10^5^ CFU/ml *M. ulcerans* JKD8049 pMV306 hsp16+luxG13. This resulted in 90% (9 out of 10) of mice presenting measurable light emission on IVIS^®^-images. Other than at the injection site at the tail, no other foci of infection as indicated by bioluminescence were observed (Fig 1B). Fifty percent (5 out of 10) gradually developed macroscopically apparent lesions resembling Buruli ulcer within 17 weeks (Fig 1C). None of the animals showed other signs of illness than skin lesion that were restricted to the approximate sites of injection.

### Course of the infection as measured by bioluminescence

To study the course of the infection in terms of bacterial burden measured in bioluminescence, mice were imaged weekly with the IVIS^®^ system. Bioluminescence, measured in emitted photons/s rose exponentially to a maximum of 1×10^7^ in week seven (Fig 1D), according to our standard curve, this equals about 5×10^6^ CFU/ml (Fig 1A) and was associated with advanced, severe pathology (Table 1). From this time-point on, the signal declined to a 1×10^5^ (corresponding to 4×10^4^ bacteria) threshold until the end of the experiment. At week eight, three mice reached humane endpoint and were culled. In examining the antibody titer levels against MU WCL, these began to rise in week 8, correlating with a decrease in photon/s counts. Animals displaying *severe* symptoms had higher photons/s counts, indicating higher bacterial burden, compared to those with *moderate* or no pathology. Photons/sec counts increased per week until week 6-8 when the infection seemed to plateau.

**Table 1:**
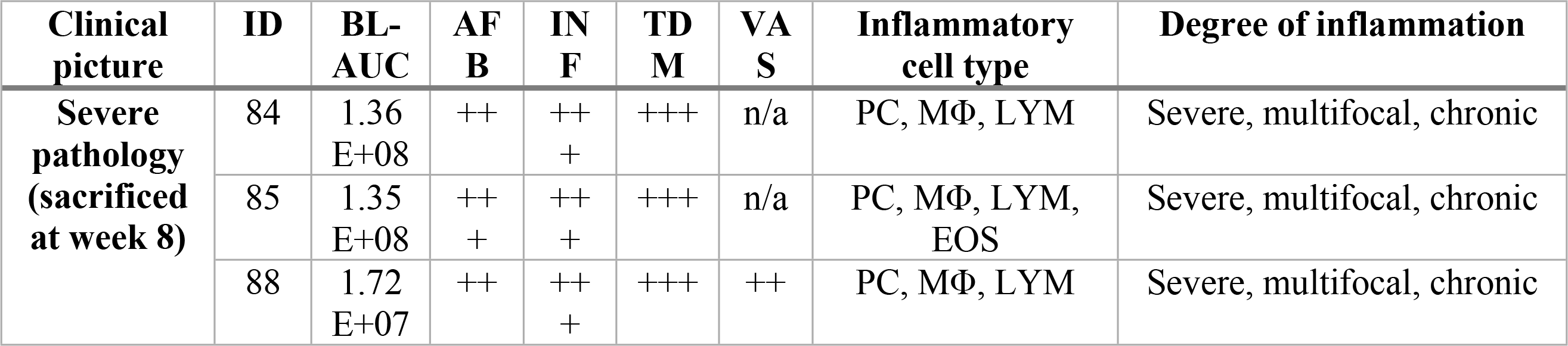
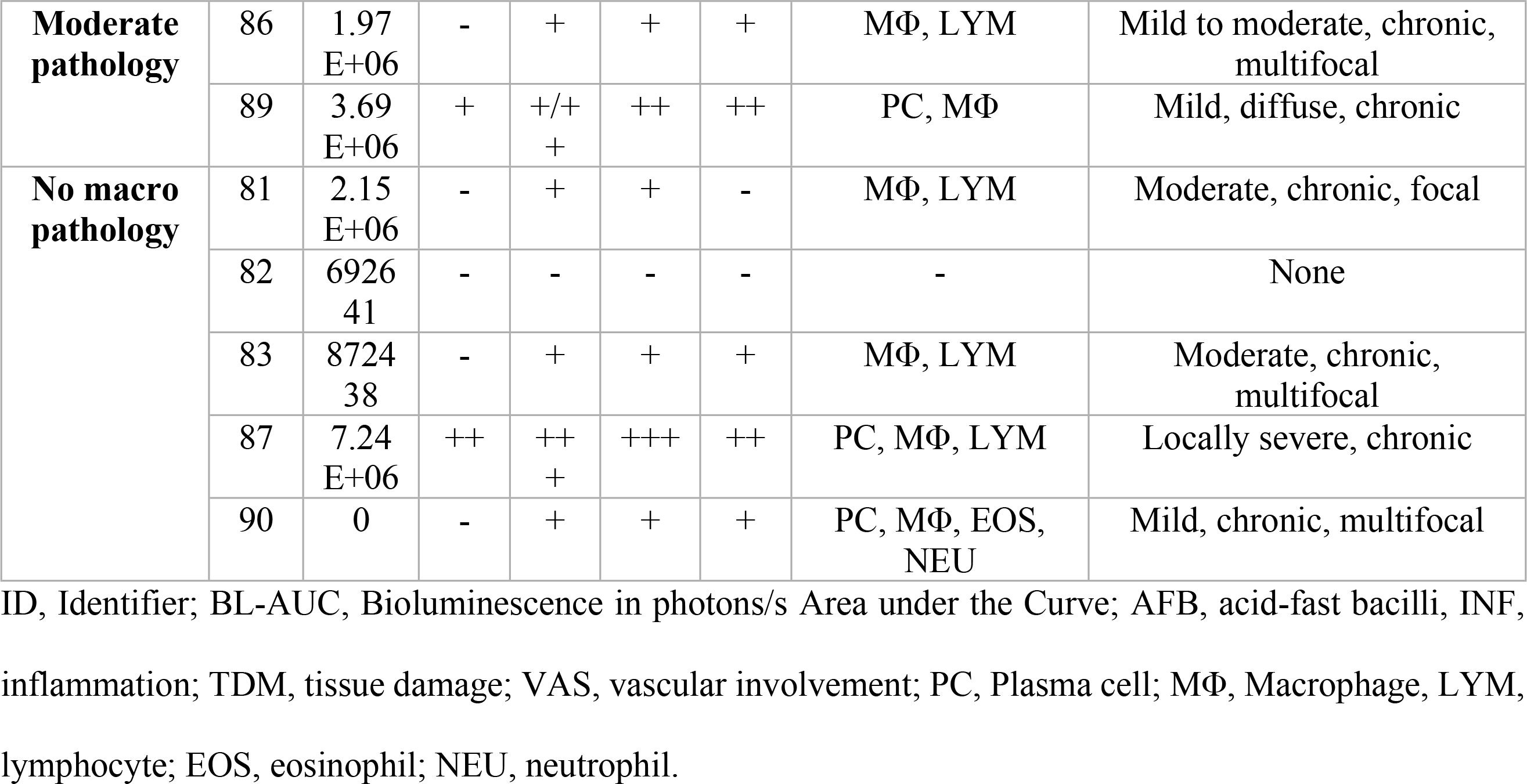
Overview of histopathological findings of mice subcutaneously infected with autoluminescent *M. ulcerans* into the tail. Animals were divided by clinical pathology in severe pathology, moderate pathology and no marcopathology. Photons per second analyzed by IVIS-imaging are shown in comparison to histological results. The amount of photos/s as a proxy for bacterial quantity correlated with pathology except in mouse 87, where no clinical pathology was seen.

### Antibody titers

To characterize the antibody-mediated immune response to *M. ulcerans*, we obtained plasma samples for ELISA at weeks 2, 4, 7, 13 and 17 of the experiment. Over time, there was a slight increase of antibody titers in response to *M. ulcerans* WCL but overall a late onset of the antibody response was noted (Fig 2A). Antibody titers reached higher levels between week 11 and 17, but overall titers, were low (Fig 2A). The response to *M. ulcerans* Hsp18 and WCL was compared and no statistically significant difference (p > 0.05) was found (Fig 2B). Furthermore, ELISA results in response to WCL at week 8 were compared between animals with severe, moderate and no clinical pathology and no statistically significant difference (p > 0.05) was found (Fig 2C,D).

**Figure 2.**
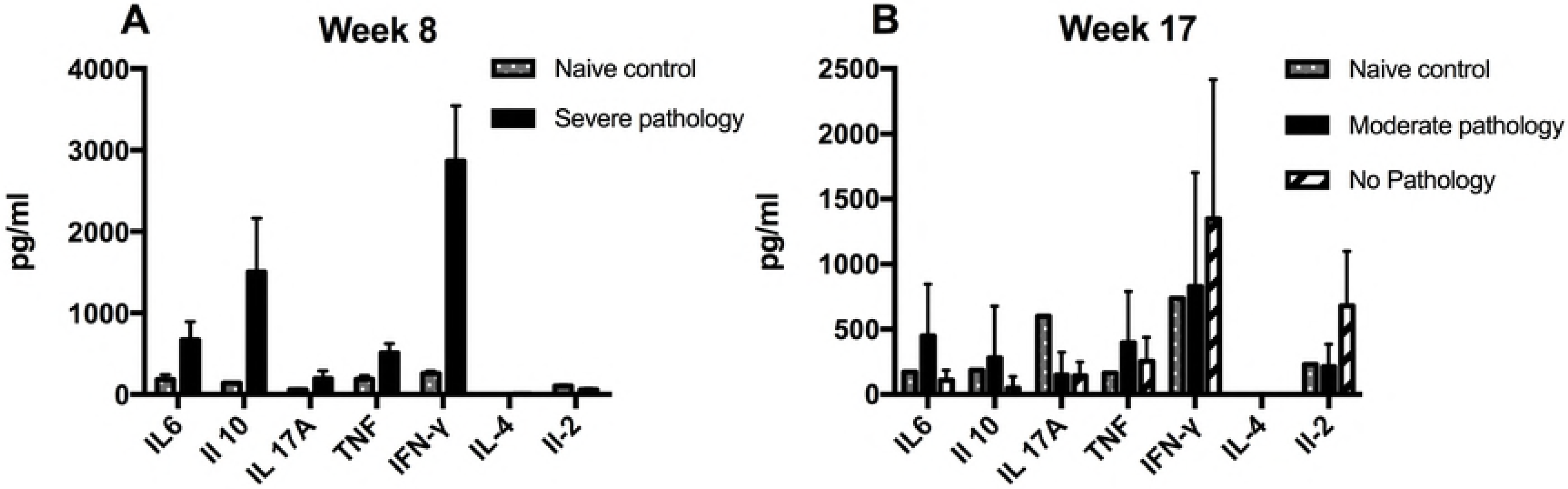
Evolution of the antibody titer against *M. ulcerans* whole cell lysate (WCL) and small heat shock protein 18 (Hsp18) measured in plasma from mice infected with *M. ulcerans* over time. A late onset of overall antibody response against an unspecific *M. ulcerans* whole cell lysate was noted in the bioluminescent *M. ulcerans* tail infection model. Antibody titers rose late, after 8 weeks and plateaued at week 13 (A). No difference (p > 0.5) was observed in the antibody titer against small heat-shock protein 18 (Hsp 18) and whole cell lysate (B). No statistically significance in antibody levels was seen between animals with severe, moderate or no apparent pathology (p > 0.5; C).

#### Late suppression of cytokines

To characterize and study the cytokine profile in our murine *M. ulcerans* infection model, intracellular cytokine staining was performed on spleen samples after eight and 17 weeks of the experiment, when three mice were humanely killed. The cytokine concentrations in splenocyte samples re-stimulated with 2μg *M. ulcerans* JKD8049 WCL were compared between mice with pathology culled at week 8 and those with and without pathology culled at week 17 were compared. In the three mice that were culled prematurely due to rapidly extending disease in week 8, elevated levels of IFN-γ and IL-10 were measured. Overall, cytokine levels were very low for all assayed cytokines in week 17 regardless of clinical state of the animal (Fig 3).

**Figure 3:**
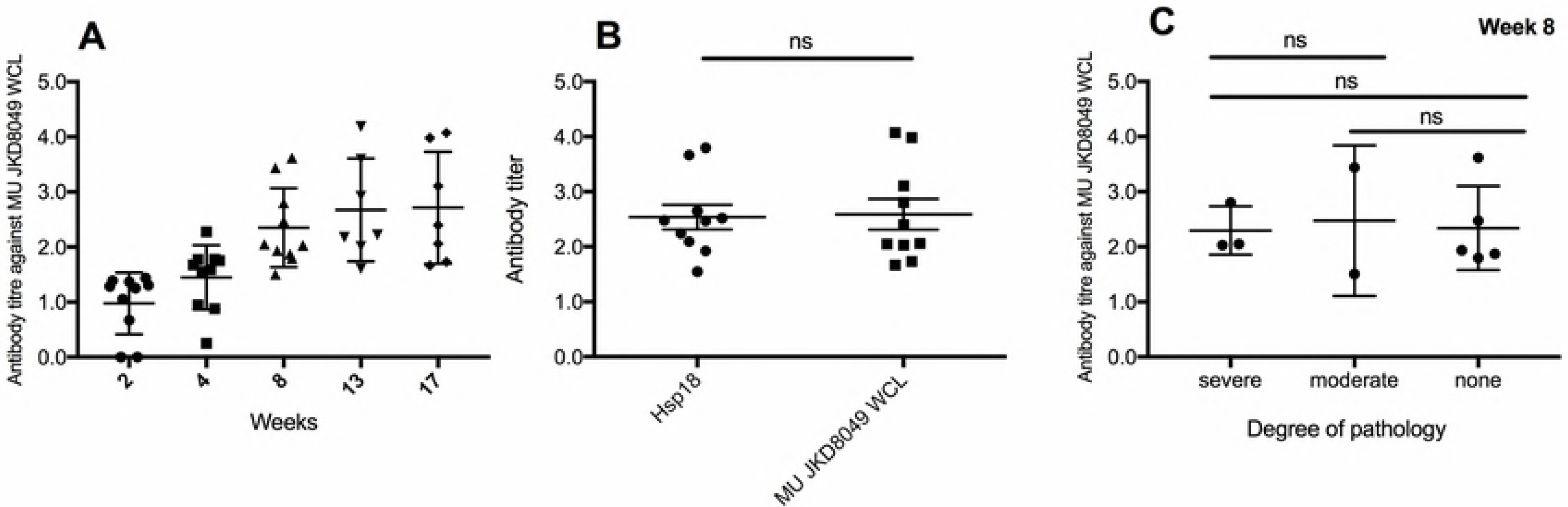
Comparison of cytokine profile assessed by intracellular cytokine staining (ICS) of mice infected with *M. ulcerans* after 8 (A) and 17 (B) weeks of infection to naïve, non-infected mice. Error bars represent standard error of the mean. Mice with advanced clinical pathology sacrificed at week eight of the experiment displayed highly elevated IFN-γ, as well as IL-6 and IL-10 levels (A). IFN-γ is known to activate macrophages and is a key regulator in granuloma formation in mycobacterial infections. At week 17, infected mice with no apparent pathology had higher IFN-γ counts than other mice, as well as slightly elevated IL-2 levels.

#### Histopathology of lesions

In order to validate the model and study the pathology of *M. ulcerans*, histopathology was performed on skin lesions and compared to those of humans described in the literature. Specimens from infected tissue were subjected to histopathological analysis in Ziehl-Neelsen and H&E-staining. Aggregates of acid-fast bacilli, *M. ulcerans*, were observed at 300 – 400 μm beneath the epidermis (Fig 4A). Furthermore, epidermal hyperplasia and immune cell infiltrates were apparent (Fig. 4). Bioluminescence (photons/s) as proxy for bacterial quantity correlated well with the histological extent of disease except for mouse ID 87 (Table 1). Numerous acid-fast bacilli as well as severe, multifocal, chronic inflammation marked by presence of plasma cells, macrophages and lymphocytes were observed in mice with severe clinical pathology. There was extensive tissue damage, as well as vascular involvement in these animals. Mice with moderate clinical pathology exhibited mild and rather diffuse histological pictures and less tissue damage. Mice that had no obvious clinical signs of disease had low bioluminescence and showed moderate to little localized/focal histological features of inflammation (Table 1, Fig 4).

**Figure 4:**
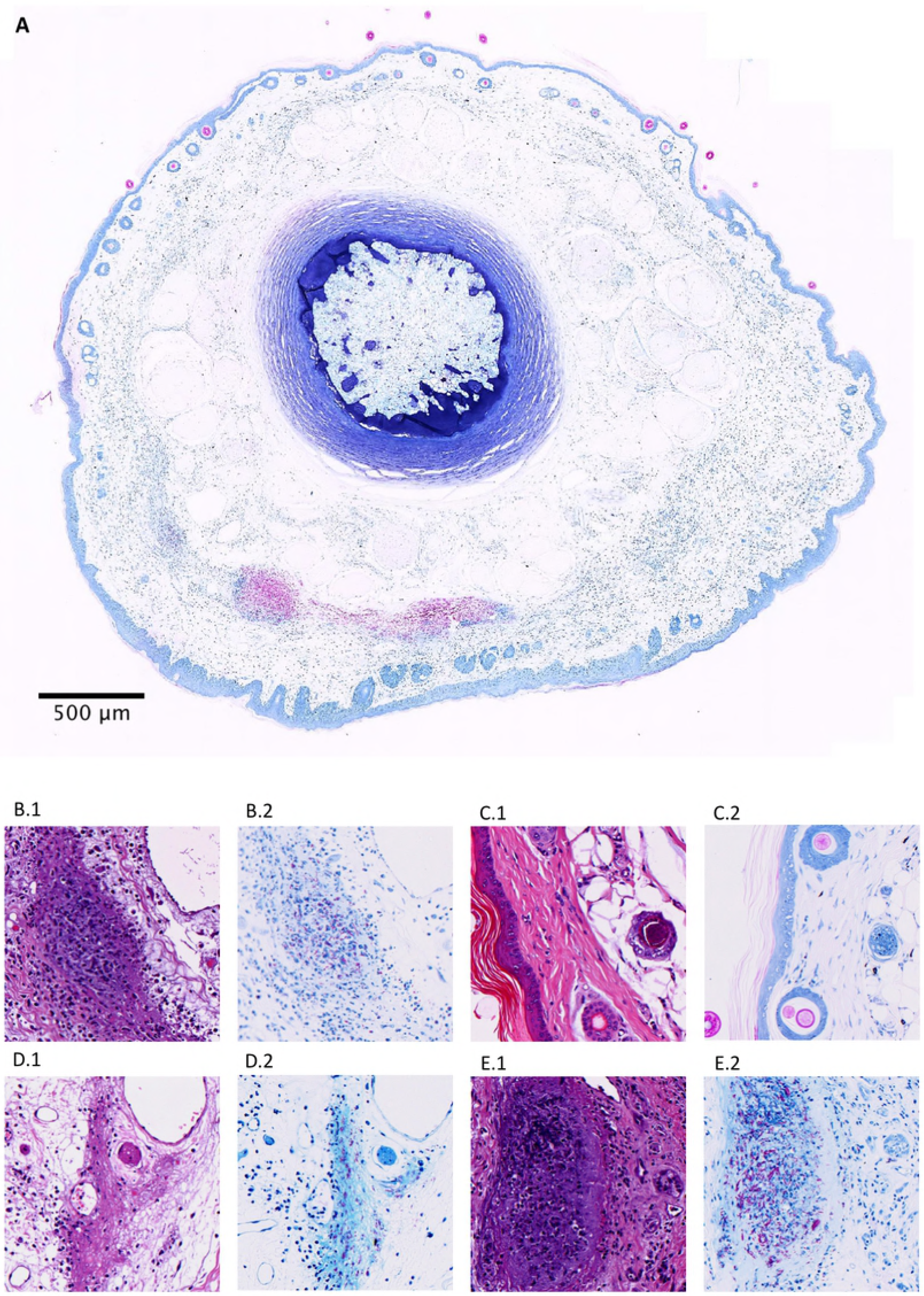
Histological specimens of mice infected with *M. ulcerans* into the tail in Hematoxylin and Eosin (H & E; A1,B1,C1,D1) or Ziel-Neelsen (ZN; A2,B2, C2,D2) staining. (A) Whole slide cross-section of mouse tail (ID #85) infected with *M. ulcerans* subcutaneously, humanely killed eight weeks’ post-infection due to advanced clinical pathology. Visible, are clusters of acid-fast bacilli and in the cutis and subcutis, approx. 300-400 μm beneath the surface. (B.1 and B.2) Example of presence of AFB in mouse # 85, as well as granuloma formation. (C.1 and C2.) Normal mouse tail histology of a naïve, uninfected mouse with thin epidermis and intact hair follicles. (D.1 and D.2) Moderate pathology (mouse #89) with diffuse inflammation, tissue damage and presence of AFB. (E1,E2) Necrosis, granuloma formation, inflammation and abundant extracellular clustering of AFB was observed in mice with severe pathology (mouse #84).

### Discussion

In this study we have successfully applied the use of bioluminescent *M. ulcerans* JKD8049 to develop a model that allows us to follow the immune response to *M. ulcerans* in BALB/c mice correlating with human Buruli ulcer disease. The onset of clinical signs was gradual and mice developed characteristic, localized lesions. Necrosis of the subcutis, chronic inflammation with the presence of macrophages and lymphocytes, granuloma formation, panniculitis as well as the and presence of AFBs correlating with disease progression are hallmarks of Buruli ulcer histopathology described in humans [49,50]. The extent and the overall of pathology observed in the mouse tail tissue in our study was comparable to the above-mentioned experience from human patients supporting the use of this mouse model to study Buruli ulcer.

The bioluminescent read-out correlated with the histopathological outcome in a dose-dependent manner, where an elevated photons/s counts, indicating high bacterial burden coincided with more progressive histological disease (Table 1) underlining the usefulness of IVIS^®^-imaging and bioluminescence as a marker for disease progression. The use of bioluminescent *M. ulcerans* permitted us to both verify the location of the bacteria as well as the growth rate in the lesion. After subcutaneous injection we observed presence of photon-emitting bacteria exclusively in the upper-third of the mouse tail, which was the site of injection.

We noticed a decline and plateauing of bioluminescence from week 8 onwards. This phenomenon could be explained by a plateauing of the bacterial growth curve in the lesion and transition into a stationary phase where less of the immunosuppressive toxin mycolactone is produced and partial host control sets in which is reflected to some extent by the rise of antibody titers around that time-point (Fig 2). The later phase of Buruli ulcer infection is characterized by granuloma formation and highly localized extracellular AFBs within these lesions. Also, a less active metabolic state during this phase could lead to decreased bioluminescence. Vasculopathy is a feature of Buruli ulcers observed in humans [49] and mice (Table 1, Fig 4). A hypoxic state within the lesion might also decrease bioluminescence and more research is needed to elucidate the usefulness of bioluminescence as a marker of bacterial quantity beyond approx. 8 weeks of infection in the BALB/c mouse.

### Immunology and course of disease

The immune response to *M. ulcerans* is influenced by the microbes’ toxin, mycolactone. Dendritic cells (DC’s) are inhibited by ML which can impair their ability to prime a cellular immune responses and phagocytose the bacteria [51]. Also, suppression of a CD4+ immune response was observed in humans [52,53] and efficient mounting of a Th1 response and elevated IFN-γ seemed protective [54]. T-cells depletion, mediated by miRNAs has also been attributed to ML {GueninMace:2011bp}. It is believed that, like in tuberculosis, an effective cell-mediated immune responses can naturally control the infection a nd are likely also important for conferring transient protection against BU, experimentally [55]. Markedly elevated cytokines in human Buruli cases are IFN-γ and IL-10 [56]. IFN-γ is known to be an early mediator of host response to *M. ulcerans* [57] and increases in patients after 4 – 8 weeks of antimicrobial treatment indicating immunocompetence against *M. ulcerans* and the mounting of a supportive CD4+ Th1-response [56]. The elevated IFN-γ response seen in mice with severe pathology (Fig 3A) can thus be interpreted as an early reaction to a large amount of actively multiplying bacteria, whereas at week 17, mice with no pathology had higher IFN-γ levels than those with pathology possibly due to sufficient host control of the pathogen. Consistently, patients with pre-ulcerative lesions (early-phase) and patients with healed lesions (host control) both showed elevated IFN-γ levels [58] whilst IL-10 seems to be somewhat nonspecifically elevated during all phases of Buruli ulcer disease [54,56,58]

In our experiment, we noted a late onset of cell-mediated immunity in the mice infected with *M. ulcerans.* Antibody titres slowly rose, but only at week 7. Interestingly, the decline in photons/s and the increase in antibody titers coincided in week 7. It is not clear if rising antibody levels helped to gain control of the infection or if declining bacterial load resulted in less immune suppression by ML and led to a reactivation of the immune system and an increase of antibody levels. In humans, serological screening for Hsp18 antibodies indicates that large parts of the population in endemic areas are exposed to *M. ulcerans* but only some develop the disease [59]. Guinea pigs infected with *M. ulcerans* appear to self-heal as do some mice [60]. It is conceivable that humans infected with certain doses of *M. ulcerans* develop either no disease, limited disease, or even unnoticed disease that self-resolves. Evidence for these scenarios has been observed in Buruli ulcer patients who have defaulted from antibiotic treatment regimens, yet could still be contacted for follow-up and showed to have healed lesions despite incomplete treatment [61]. It is likely, that the bacterial burden was not zero in these patients at the time of default but that it reached a critical nadir at which host immunity overcame the counteracting effect of mycolactone and controlled infection. Individuals that are able to establish an efficient immune response to MU might control and clear the infection unnoticed as was the case with 50% of subcutaneously infected mice in our experiment. We have previously deduced a low infectious dose 50% (ID50) of <10 CFU from experiments involving mechanical injury simulated by needle stick to *M. ulcerans* externally contaminated mouse tails. Even though in this current research bacterial presence was measured by IVIS^®^ imaging in 90% of mice, only 50% of animals showed clinical disease in this experiment following subcutaneous injection with approx. 5×10^6^ CFU/ml. This observation and discrepancy with our previous research might be explained by the different handling of the inoculating needle, perpendicular, and supposedly deeper penetration in the study by Wallace et al. [9] and more superficial penetration in a 20-30° angle in subcutaneous injections in this study. Experiments are required to assess the *M. ulcerans* ID50 using carefully controlled inoculation conditions, perhaps using a micromanipulator with decreasing doses of *M. ulcerans*.

At week 8, three animals of the SC cohort had reached the humane experimental endpoint and were culled. Their cytokine profile data were comparable to those of the other animals culled at week 17 and provide an insight into the evolution of the cytokine profile. While there were considerable amounts of IFN-γ and IL-10 indicating a Th1-mediated response observed in the animals sacrificed at week 8, all cytokine levels were reduced by week 17. The suppression of cytokines is due to inhibition by ML of nascent membrane and secretory proteins egress through the ER membrane [6,62]. In human patients with Buruli ulcer, overall suppressed IFN-γ levels are seen [54]. The triad of peak bacterial load with worsening pathology, low but rising antibody titers and elevated Th1 subset cytokines in week 7 could also explain the paradoxical response seen in patients; an overreaction and inflammation of the reactivating immune system noted in patients beginning antibiotics for Buruli ulcer [63]. Animals not showing clinical signs of disease had marginally higher antibodies levels against *M. ulcerans* (Fig 2) possibly indicating some sort of immune response to the infection, though the results were statistically not significant in our small sample.

We demonstrated virulence of the *M. ulcerans*+pMV306 hsp16+luxG13 reporter strain and observed localized clinical disease with human-like pathology in 50% of the animals after inoculation with 5×10^6^ CFU/ml bacteria. IVIS^®^-imaging of *M. ulcerans* infection can reduce and refine animal usage in Buruli ulcer research and enables further studies of into pathology of the disease as well as pre-clinical drug and vaccine evaluation.

## Acknowledgements

We thank Rolfe Howlett and John Hayman and for help with analyzing and scoring the histopathological results.

